# First-in-Class Small Molecule ROBO2 Binders Identified through Integrated Virtual Screening and Biophysical Validation

**DOI:** 10.64898/2026.01.05.697640

**Authors:** Kirti Upmanyu, Hossam Nada, Moustafa T. Gabr

## Abstract

Roundabout homolog 2 (ROBO2) is a transmembrane receptor implicated in glioblastoma progression through its interaction with Slit2-mediated signaling pathways. Dysregulated Slit2–ROBO2 signaling enhances tumor cell migration, invasion, and tissue infiltration, while elevated ROBO2 levels contribute to an immunosuppressive tumor microenvironment supporting GBM aggressiveness and highlighting ROBO2 as a therapeutic target. Despite its therapeutic relevance, no ROBO2-targeted small molecules have been reported. To address this gap, we performed a structure-based virtual screening campaign targeting ROBO2, followed by experimental validation with Dianthus TRIC platform and microscale thermophoresis (MST). Fifteen compounds were screened for ROBO2 binding, from which four candidates exhibited robust TRIC signals. Subsequent affinity measurements revealed that two small molecules, Z1334432986 and Z1692774161, bind ROBO2 in a reproducible concentration-dependent manner, with dissociation constants (Kd) of 40.8±4.8 μM and 25.8±16.95 μM, respectively. Molecular docking with validated hits revealed a shared ROBO2 binding pocket defined by conserved anchor residues (ASN354, SER366, ASP385) and accommodation of distinct ligand conformations within the ROBO2 binding pocket. This work establishes a screening pipeline for identifying ROBO2-targeted small molecules and lays the foundation for developing therapeutics aimed at disrupting Slit–ROBO2 signaling in GBM.

**Graphical abstract:** 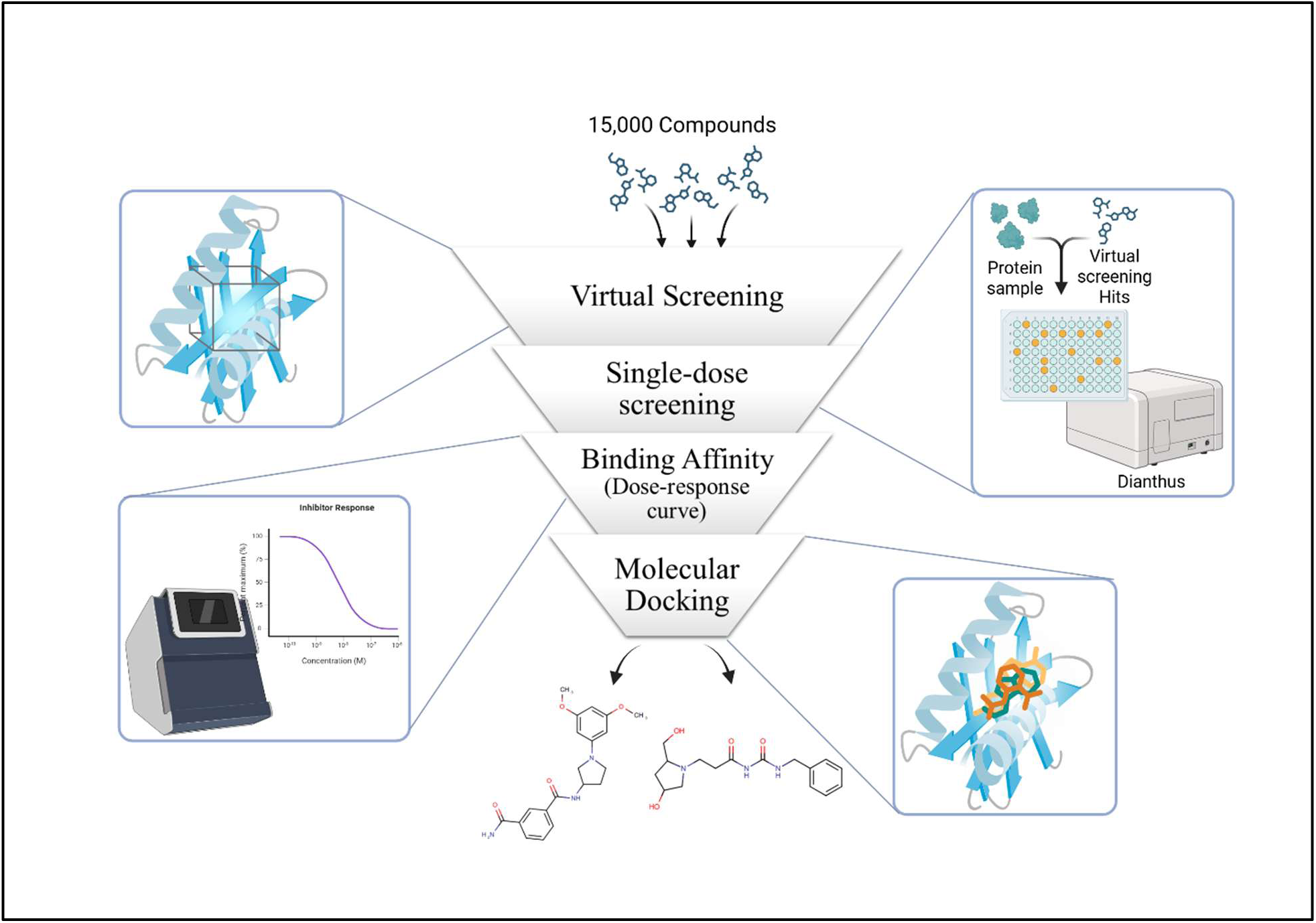

## 1. Introduction

Roundabout homolog 2 (ROBO2) is a single spanning transmembrane receptor of the immunoglobulin superfamily and plays a crucial role in axon guidance and cell migration [1, 2]. Structurally it is composed of an extracellular ectodomain containing five Ig-like domains followed by three fibronectin type-III (FNIII) repeats, a single-pass transmembrane helix, and a cytoplasmic tail with conserved CC0–CC3 intracellular motifs responsible for recruiting downstream signaling partners [3, 4]. Through this architecture ROBO2 recognize extracellular signals and convert them into cytoskeletal and transcriptional responses that shape neuronal connectivity [5, 6].

ROBO2 primarily interacts with Slit2, a large, secreted glycoprotein that undergoes proteolytic cleavage into N-terminal (Slit2-N) and C-terminal fragments [7–9]. The N fragment of Slit2 binds with high affinity to the first Ig domain of ROBO2, triggering receptor clustering and activation of intracellular signaling cascades that regulate cell migration, cytoskeletal dynamics, inflammatory responses, growth-cone repulsion, and axon navigation [10]. Beyond development, Slit2– ROBO2 signaling modulates synaptic plasticity, dendritic spine remodeling, and blood–brain barrier integrity [11].

Increasing evidence indicates that ROBO2 plays context-dependent roles in cancer, functioning as either a suppressor or modulator of tumor–stromal interactions [12]. Recent studies have identified the Slit–Robo signaling axis as a key pathway that may regulate glioblastoma invasiveness [12]. Among Robo family members encoded by mammals (ROBO1-4), ROBO2 particularly showed strong clinical and experimental associations with aggressive disease [13]. Analysis of the NCI REMBRANDT dataset reveals that glioma patients with more than twofold ROBO2 overexpression experience significantly reduced overall survival (17.1 vs. 37.4 months), suggesting a correlation between elevated ROBO2 and poor prognosis [14]. Its overexpression, association with poorer survival, and functional relevance to tumor biology position ROBO2 as a promising therapeutic target within the Slit–ROBO pathway for counteracting glioblastoma [15]. Functionally, siRNA-mediated silencing of ROBO2 in U87 glioma cells showed a measurable survival advantage in vivo and supporting a role for ROBO2 in promoting tumor aggressiveness [14].

Despite growing evidence that the Slit– ROBO2 signaling pathway plays a critical role in tumor progression, immune modulation, and fibrotic processes, there are currently no approved or well-characterized small-molecule inhibitors targeting the Slit– ROBO2 (or Slit– ROBO1) interaction for therapeutic use in cancer or fibrosis. This represents a significant gap in drug development, particularly given the pathway’s involvement in promoting tumor aggressiveness and shaping the tumor microenvironment. Biologic-based strategies, such as monoclonal antibodies or soluble receptor decoys, have demonstrated proof-of-concept for inhibiting Slit–Robo signaling [16, 17]. However, these approaches face substantial translational challenges in central nervous system (CNS) malignancies such as glioblastoma (GBM). The blood–brain barrier (BBB) severely limits the delivery of large protein therapeutics, while poor tissue diffusion and potential immunogenicity further restrict their clinical utility [18].

In this context, small-molecule inhibitors offer a promising alternative. Their superior tissue penetration, manageable pharmacokinetics, and lower immunogenicity make them particularly well-suited to targeting CNS tumors [19, 20]. By disrupting the Slit– ROBO2 axis, these compounds could effectively “reprogram” the tumor microenvironment, reduce immunosuppressive signaling, and enhance the efficacy of immunotherapy or chemotherapy. Developing such inhibitors would not only address a critical unmet need in GBM therapy but also provide a broadly applicable strategy for targeting Slit–ROBO signaling in other malignancies and fibrotic diseases.

In this study, we aimed to target the critical Slit2– ROBO2 signaling axis by developing a robust pipeline for identifying small-molecule modulators of ROBO2. Using structure-based virtual screening, we computationally screened a library of 15,000 compounds and prioritized fifteen candidate compounds, which were then experimentally evaluated using the Dianthus platform. Subsequently dose-dependent validation was performed with microscale thermophoresis (MST) to determine their binding affinities. ultimately identifying two promising small molecules, Z1334432986 and Z1692774161, that reproducibly bind ROBO2. By developing a validated workflow for discovering and characterizing ROBO2-targeting compounds, this study provides the first small-molecule modulators of ROBO2 highlighting new possibilities for therapeutically interfering with Slit2– ROBO2 signaling. These compounds could potentially inhibit glioblastoma cell migration and invasion, reshape the tumor microenvironment, and provide a strategy for targeting ROBO2-driven diseases.

## 2. Methods

### 2.1 Virtual Screening

The crystal structure of human ROBO2 extracellular domains 4-5 (PDB ID: 5NOI) was retrieved from the Protein Data Bank and used as the target structure for virtual screening [21]. This structure encompasses two immunoglobulin-like domains that play critical roles in SLIT ligand recognition and receptor dimerization. The structure was prepared for docking studies using the Protein Preparation Wizard in Schrödinger Maestro, which included addition of hydrogen atoms, assignment of appropriate protonation states at pH 7.4, optimization of hydrogen bond networks, and restrained energy minimization using the OPLS4 force field. The prepared ROBO2 structure was analyzed using PrankWeb, a web-based platform for ligand binding site prediction. PrankWeb utilizes machine learning algorithms to identify potential binding pockets based on structural features, evolutionary conservation, and geometric properties [22].

Structure-based virtual screening was performed using the Maestro Schrödinger suite. An in-house chemical library containing 15,000 Enamine compounds was screened against the identified ROBO2 binding pocket (PDB ID: 5NOI. The virtual screening workflow consisted of multiple stages:

1. Ligand Preparation: All 15,000 compounds from the Enamine library were prepared using LigPrep module, which generated tautomeric and ionization states at pH 7.0 ± 2.0, optimized geometries, and generated low-energy 3D conformers.
2. Receptor Grid Generation: A docking grid was centered on the predicted binding pocket identified by PrankWeb, with the inner box dimensions set to capture the entire pocket region and the outer box providing adequate space for ligand sampling.
3. Hierarchical Docking Protocol: The virtual screening workflow employed a multi-stage filtering approach. Initial high-throughput virtual screening (HTVS) was performed on all 15,000 compounds using simplified scoring to rapidly eliminate non-binders. The top-ranked compounds progressed to standard precision (SP) docking for more accurate pose prediction and scoring. Finally, the highest-scoring candidates underwent extra precision (XP) docking with enhanced sampling and a more rigorous scoring function to identify the most promising hits.

### 2.2 Single dose Screening using Dianthus

The compounds identified by virtual screening were initially analyzed for their binding with ROBO2 using microscale thermophoresis. Briefly, 50nM of His-tag ROBO2 was mixed with 25nM dye in phosphate buffer saline supplemented with 0.05% Tween-20 (pH 7.4) and incubated for 30 minutes at room temperature in dark. Finally, the assay comprised of 25 nM of protein and 100 μM of compound in PBST. Samples were allowed to incubate for 30 minutes at room temperature, then briefly spun down before being assessed on a Dianthus NT.23 Pico instrument. This system has been robustly validated in our laboratory for similarly challenging protein–ligand interactions, including the immune co-stimulatory receptor CD28 and the axon guidance ligand SLIT2 [23, 24]. Buffer containing only DMSO served as the negative control. Each experiment was conducted in triplicate, and the values reported represent the mean of those replicates.

The inherent fluorescent property of compounds was analyzed by running control reactions with only 100 μM of each compound in PBST in the absence of dye and protein and incubated under identical conditions as above. Furthermore, secondary check for the compound’s potential to diminish or ‘quench’ the overall fluorescent signal of the experimental dyes was assessed by mixing each test compound with the 25 nM RED-tris-NTA, and subsequently monitoring the resultant signal intensity.

### 2.3 Dose Response using Monolith

The compounds that showed a clear binding signal in the single dose screening were further followed up using dose-response MST to determine their binding affinity with ROBO2. A sixteen-point serial dilution series was prepared beginning at 250 μM in PBST extending into the low-nanomolar range, and each dilution was combined with 25nM fluorescently labeled His-tag ROBO2 protein while keeping the final DMSO concentration at 2%. After equilibrating the mixtures for 30 minutes at room temperature, samples were transferred into standard MST capillaries and analyzed on a Monolith NT.115 using medium–high infrared laser settings and 60–80% LED excitation in the red channel. This MST platform has been extensively validated in our laboratory for high-affinity and low-affinity interactions involving difficult targets, including the immune co-stimulatory receptor CD28 and the pseudoenzymatic immune checkpoint CHI3L1 [25, 26]. Dissociation constants (Kd values) were determined with the MO.Affinity Analysis software (NanoTemper Technologies).

### 2.4. Molecular docking visualization

The docking poses and protein-ligand interactions of the experimentally validated hits were analyzed and visualized using BIOVIA Discovery Studio Visualizer (Dassault Systèmes). Two-dimensional ligand interaction diagrams were generated to map hydrogen bonds, hydrophobic interactions, π-π stacking, π-sulfur interactions, and other non-covalent contacts between the ligands and ROBO2 residues. Hydrogen bonds were identified using distance cutoffs of ≤3.5 Å between donor and acceptor heavy atoms and angle cutoffs of ≥120° for donor-H-acceptor angles. Hydrophobic interactions were defined as contacts between hydrophobic residues and ligand carbon atoms within 4.0 Å. All molecular graphics were prepared using Discovery Studio Visualizer with standard atom coloring (carbon in gray, nitrogen in blue, oxygen in red, sulfur in yellow) and interaction indicators (hydrogen bonds as green dashed lines, hydrophobic contacts as pink/purple circles).

## 3. Results

### 3.1 Virtual screening

The ROBO2 protein structure was analyzed using PrankWeb, a web-based platform for ligand binding site prediction. PrankWeb utilizes machine learning algorithms to identify potential binding pockets based on structural features and conservation scores. The analysis identified a high-confidence binding pocket with a score of 2.10 and probability of 0.047 (Figure SX), comprising 10 residues. The predicted pocket showed an average conservation score of 1.173 where the binding site residues were distributed across a conserved leucine-rich repeat region consistent with the immunoglobulin-like domain 1 (Ig1) of ROBO2.

The virtual screening identified 15 potential hits with favorable predicted binding affinities to the ROBO2 pocket. The docking scores ranged from −7.1 kcal/mol (compound Z1177048571, the top hit) to −6.037 kcal/mol (compound Z595692196). All selected compounds demonstrated docking scores below −6.0 kcal/mol, suggesting reasonable predicted binding affinity. The distribution of scores showed a relatively narrow range of approximately 1 kcal/mol between the best and 15th-ranked compounds, indicating a cluster of similarly ranked candidates rather than a single clear outlier.

**Table 1.** Virtual screening results.

| Compound ID | Docking score (kcal/mol) |
| --- | --- |
| Z1177048571 | -7.1 |
| Z1692774161 | -6.879 |
| Z2067638674 | -6.801 |
| Z743902858 | -6.731 |
| Z1839977548 | -6.629 |
| Z2376885317 | -6.477 |
| Z1121000042 | -6.47 |
| Z1334432986 | -6.438 |
| Z1246861296 | -6.21 |
| Z3476327031 | -6.192 |
| Z5860376961 | -6.173 |
| Z2755578126 | -6.135 |
| Z1416424828 | -6.106 |
| Z595692204 | -6.083 |
| Z595692196 | -6.037 |

### 3.2 Initial screening identified four ROBO2 binding compounds with Dianthus

Compounds identified as hits from the virtual screening were further evaluated for their binding to ROBO2 using a single-concentration assay at 100 μM on the Dianthus platform, utilizing Temperature-Related Intensity Change (TRIC) with a 10-second measurement interval. To determine significant binding, compounds were considered positive if their signal exceeded five times the standard deviation of the negative control, which consisted of protein and dye alone. Out of the 15 compounds tested, four compounds, exhibited reproducible and robust binding to ROBO2.

The compounds were also analyzed for the inherent fluorescence that could interfere with the signal by comparing the fluorescence signal generated by each compound with the control containing only 2% DMSO. None of the tested compounds showed auto fluorescence signals when compared to the control. Similarly, the compounds were also analyzed for their ability to quence the fluorescence signal and only 1 compound, Z1416424828 showed the quenching ability and was not included further. The initial screening gave four ROBO2 binding compounds, Z1334432986, Z1692774161, Z1177048571, Z1839977548 which were further evaluated for dose-dependent response using MST.

### 3.3 Binding Affinity of ROBO2 and Slit-2 using MST

To establish optimal MST conditions for the binding assay of ROBO2 and small molecules we first characterized the binding of ROBO2 with its native ligand Slit2. The twelve-point serial dilution series of Fc tagged Slit2 was set up ranging from 1 μM to lower nanomolar concentration. Each concentration was titrated with 25 nM of fluorescently labeled His Tag ROBO2 in PBST along with 2% DMSO. The resulting response evaluation was done with the spectral shifts (Ratio 670/650) displaying a clear, concentration-dependent response, allowing robust curve fitting using a standard Kd model. This analysis yielded a dissociation constant of 18.9±8.25 nM (Figure 2A), confirming strong and specific Slit2– ROBO2 binding under these conditions and establishing a reliable MST assay framework for subsequent evaluation of small-molecule hits.

**Figure 1:**
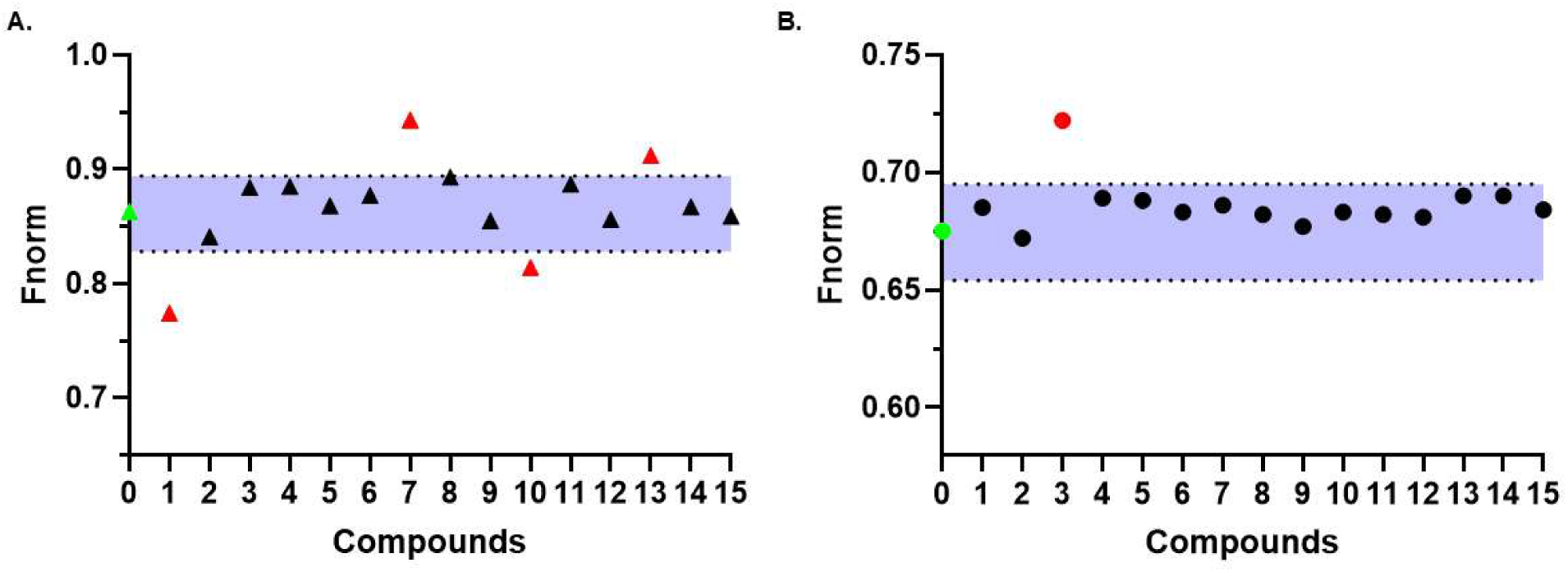
Single-dose screening of virtual-screening hits for ROBO2 binding using Dianthus. (A) Primary screening of 15 compounds for ROBO2 binding identified four potential compounds (Z1334432986, Z1692774161, Z1177048571, Z1839977548) represented with red dots and the black dots represent the non-binders (Compounds exceeding five times the standard deviation of the negative control, which included protein and dye in 2% DMSO, were classified as binders). (B) Compounds were assessed for auto-fluorescence and fluorescence quenching relative to a 2% DMSO control; none displayed auto-fluorescence, and only one compound (Z1416424828) showed quenching and was excluded.

**Figure 2:**
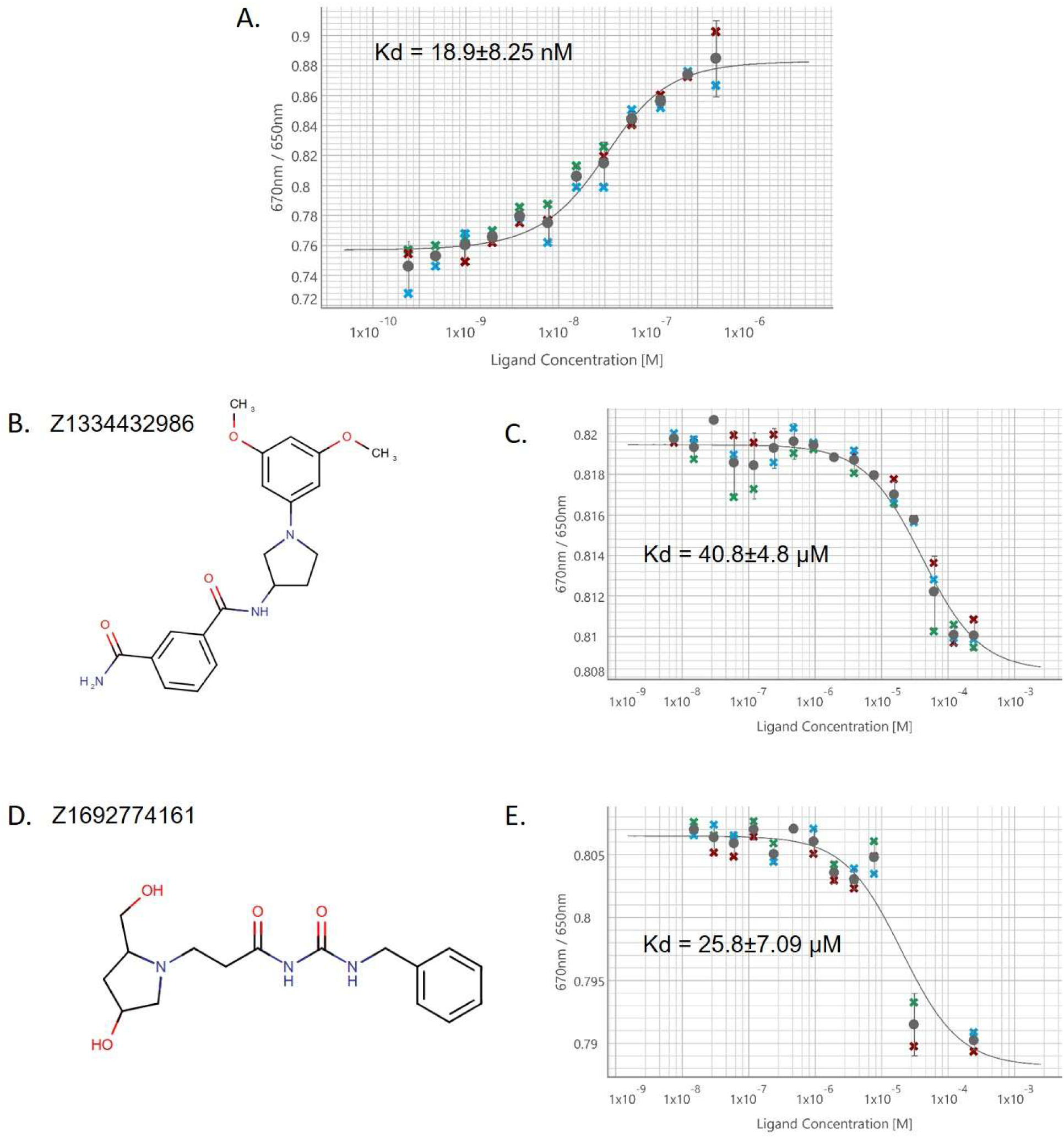
MST-based binding analysis of Slit2–ROBO2 and ROBO2-binding small molecules. (A) Dose-response binding curve of serially diluted Fc-Slit2 wi0th 25 nM His-tagged ROBO2 yielding a Kd of 18.9 ± 8.25 nM. (B) Structure of compound 1334432986. (C) Dose-response binding curve of ROBO2 with compound Z1334432986, yielding a Kd of 40.8±4.8 μM. (D) Structure of compound Z1692774161. (E) Dose-response binding curve of ROBO2 with compound Z1692774161 (n=3, error bars display SD).

### 3.4 Binding Affinity of small molecules with ROBO2

The compounds identified as hits from dianthus single-dose screening were further validated to quantitatively evaluate the direct binding interactions and determine their binding affinity to ROBO2 using MST. Serial dilutions series (250 μM to nanomolar concentrations) of each compound was prepared and titrated with 25 nM fluorescently labeled His-tag ROBO2 protein in optimized PBST buffer conditions with 2% DMSO. The resulting response evaluation with spectral shift produced well-defined binding curves, which were analyzed using a standard Kd fitting model. Of the four tested compounds, two compounds, Z1334432986 (Figure 2B) and Z1692774161 (Figure 2D) exhibited clear, reproducible, dose-dependent response, corresponding to dissociation constants (Kd) of 40.8 μM (Figure 2C) and 20.8 μM (Figure 2E), respectively. The other two compounds showed no binding or non-reproducible responses. This is the first study to report small molecules (Z1334432986 and Z1692774161) that interact with ROBO2 in a dose-dependent manner.

### 3.4. Binding Modes of the validated hits

The predicted binding modes of the two experimentally validated compounds (Z1692774161and Z1334432986) suggested the presence of key features of the ROBO2 binding pocket. Both compounds engage a common set of residues including ASN354, SER366, and ASP385 (Figure 3) which suggests that these are key anchor points for ligand binding. The consistent involvement of these residues across different chemical scaffolds indicates they likely represent hot spots within the binding site that contribute disproportionately to binding affinity. The binding pocket exhibited the ability to accommodate diverse chemical architectures where Z1692774161 adopts an extended conformation spanning the pocket length, while Z1334432986 adopts a more compact, L-shaped geometry that exploits pocket depth. This structural plasticity suggests the binding site may be amenable to further optimization through structure-activity relationship (SAR) studies exploring both linear and angular molecular geometries. Interestingly, both compounds featured terminal aromatic rings which were predicted to establish a π-π interaction with LEU356. Moreover, the hydrogen bonding networks observed in both compounds highlight the importance of properly positioned hydrogen bond donors and acceptors. This suggests that introducing additional hydroxyl or amide groups in appropriate positions could enhance binding affinity in future optimization efforts

**Figure 3:**
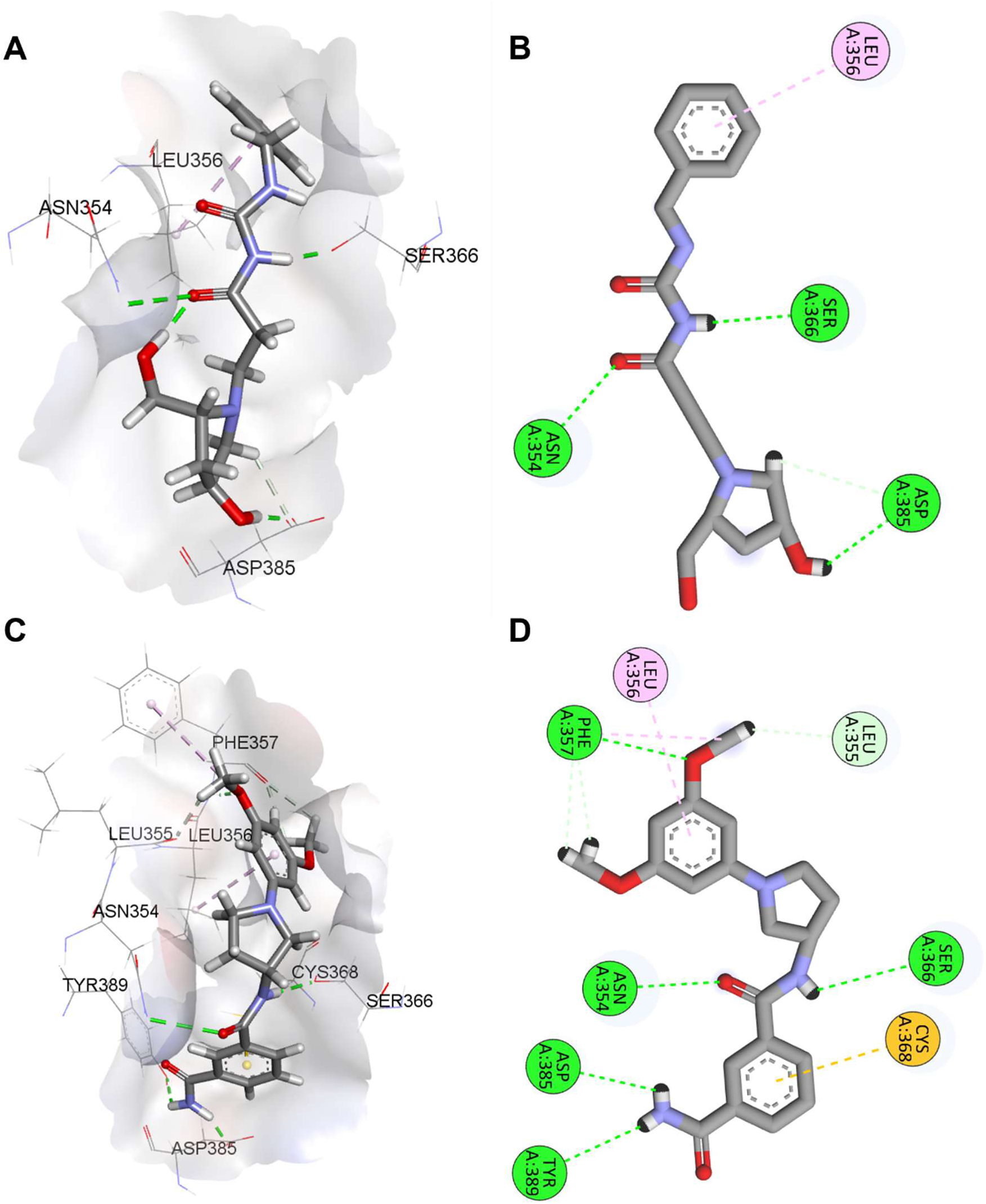
Molecular Docking Analysis of Validated Hit Compounds. A) Z1692774161 3D Binding Mode. B) Z1692774161 2D Interaction Diagram. C) Z1334432986 3D Binding Mode. Panel D) Z1334432986 2D Interaction Diagram.

## 4. Discussion

The need for new therapeutic strategies against GBM remains urgent, especially those targeting molecular pathways linked to aggressive tumor behavior and poor patient outcomes [27]. Among these, the transmembrane receptor ROBO2, which mediates Slit2-dependent guidance cues, emerges as a particularly compelling target. High ROBO2 expression is strongly associated with invasive tumor growth, an immunosuppressive microenvironment, and significantly reduced survival rates in GBM patients, highlighting the clinical significance of ROBO2 [28]. Despite this, no small-molecule inhibitors have been reported to specifically disrupt the Slit2– ROBO2 interaction, leaving a critical gap in GBM therapeutic development.

To address this, we employed a systematic screening and validation pipeline that integrates structure-based virtual screening with experimental evaluation using the Dianthus TRIC platform and MST [29, 30]. ROBO2 structural analysis showed the feasibility of targeting the protein with small molecules. Binding-site prediction using PrankWeb identified a high-confidence pocket within a conserved region of the Ig1 domain, consistent with a functionally relevant ligand-binding interface. Virtual screening of 15,000 compounds identified 15 compounds (0.1%) as ROBO2 binders with a narrow distribution of docking scores among the top-ranked compounds, suggesting that the pocket accommodates multiple chemotypes with comparable predicted affinities. Further screening with single-dose response using dianthus identified four compounds (0.02%) displaying binding with ROBO2. Further, the binding affinity of these compounds was analyzed with dose response curve analysis. This approach not only confirmed the expected high-affinity binding of the native ligand Slit2, with a dissociation constant (Kd) of 18.9 ± 8.25 nM, but also provided a reliable foundation for discovering novel small-molecule modulators of ROBO2. Our most significant finding is the identification of two small molecules, Z1334432986 and Z1692774161, which bind ROBO2 in a reproducible, concentration-dependent manner. These compounds exhibited Kd values of 40.8±4.8 μM and 25.8±16.95 μM, respectively. The observed micromolar affinities demonstrated the ability of compounds for engagement with ROBO2, but further optimization will be necessary to enhance potency and achieve affinity values closer to that of the native Slit2 ligand. Binding mode analysis of the two validated hits indicated recurrent interactions with conserved residues ASN354, SER366, and ASP385, identifying these positions as key anchoring sites within the pocket. Despite engaging common residues, the ligands adopted distinct conformations, reflecting conformational flexibility of the binding site. Together with predicted π–π interactions involving LEU356 and conserved hydrogen-bonding patterns, these findings support the structural tractability of the ROBO2 pocket and provide a basis for subsequent structure–activity relationship optimization.

This work successfully establishes a robust screening and validation pipeline capable of identifying novel ROBO2 modulators. The discovery of Z1334432986 and Z1692774161 lays the foundational chemical groundwork necessary for future therapeutic development. Continued optimization of these initial hit compounds is essential for creating potent agents capable of disrupting Slit–ROBO2 signaling, potentially offering a new treatment modality for GBM and other pathologies driven by ROBO2 dysregulation.

## Acknowledgements

This work was supported by the National Cancer Institute (NCI) under grant number R01CA293456 (PI: Gabr). We would like to thank the Fisher Drug Discovery Resource Center of Rockefeller University (RRID:SCR_020985) for providing access to the Dianthus and Monolith instruments.

## CRediT authorship contribution statement

**Kirti Upmanyu:** Writing – original draft, Visualization, Methodology, Investigation, Formal analysis, Data curation. **Hossam Nada:** Writing – original draft, Methodology, Data curation. **Gabr Moustafa:** Writing – review & editing, Supervision, Funding acquisition.

## References

[1] R. Kraut and K. Zinn, “Roundabout 2 regulates migration of sensory neurons by signaling in trans, (in eng)”, Curr Biol, vol. 14, no. 15, pp. 1319–29, Aug 10 2004, doi: 10.1016/j.cub.2004.07.052.

[2] J. H. Simpson, T. Kidd, K. S. Bland, and C. S. Goodman, ”Short-Range and Long-Range Guidance by Slit and Its Robo Receptors: Robo and Robo2 Play Distinct Roles in Midline Guidance,” Neuron, vol. 28, no. 3, pp. 753–766, 2000/12/01/ 2000, doi: 10.1016/S0896-6273(00)00151-3.

[3] J. S. Pak et al., “NELL2-Robo3 complex structure reveals mechanisms of receptor activation for axon guidance,” (in eng), Nat Commun, vol. 11, no. 1, p. 1489, Mar 20 2020, doi: 10.1038/s41467-020-15211-1.

[4] N. Yamamoto et al., “Robo2 contains a cryptic binding site for neural EGFL-like (NELL) protein 1/2,” (in eng), J Biol Chem, vol. 294, no. 12, pp. 4693–4703, Mar 22 2019, doi: 10.1074/jbc.RA118.005819.

[5] N. Sanhueza, E. C. Avilés, and C. Oliva, “The Slit-Robo signalling pathway in nervous system development: a comparative perspective from vertebrates and invertebrates,” (in eng), Open Biol, vol. 15, no. 7, p. 250026, Jul 2025, doi: 10.1098/rsob.250026.

[6] M. Wurmser, M. Muppavarapu, C. M. Tait, C. Laumonnerie, L. M. González-Castrillón, and S. I. Wilson, “Robo2 Receptor Gates the Anatomical Divergence of Neurons Derived From a Common Precursor Origin,” (in eng), Front Cell Dev Biol, vol. 9, p. 668175, 2021, doi: 10.3389/fcell.2021.668175.

[7] K. L. Whitford et al., ”Regulation of Cortical Dendrite Development by Slit-Robo Interactions,” Neuron, vol. 33, no. 1, pp. 47–61, 2002/01/03/ 2002, doi: 10.1016/S0896-6273(01)00566-9.

[8] F. Bisiak and A. A. McCarthy, ”Structure and Function of Roundabout Receptors,” in Macromolecular Protein Complexes II: Structure and Function, J. R. Harris and J. Marles-Wright Eds. Cham: Springer International Publishing, 2019, pp. 291–319.

[9] N. Rama et al., “Slit2 signaling through Robo1 and Robo2 is required for retinal neovascularization,” (in eng), Nat Med, vol. 21, no. 5, pp. 483–91, May 2015, doi: 10.1038/nm.3849.

[10] M. F. Wu, C. Y. Liao, L. Y. Wang, and J. T. Chang, “The role of Slit-Robo signaling in the regulation of tissue barriers,” (in eng), Tissue Barriers, vol. 5, no. 2, p. e1331155, Apr 3 2017, doi: 10.1080/21688370.2017.1331155.

[11] Y. Gonda, T. Namba, and C. Hanashima, “Beyond Axon Guidance: Roles of Slit-Robo Signaling in Neocortical Formation,” (in eng), Front Cell Dev Biol, vol. 8, p. 607415, 2020, doi: 10.3389/fcell.2020.607415.

[12] A. V. Pinho et al., “ROBO2 is a stroma suppressor gene in the pancreas and acts via TGF-β signalling,” (in eng), Nat Commun, vol. 9, no. 1, p. 5083, Nov 30 2018, doi: 10.1038/s41467-018-07497-z.

[13] T. A. Evans, C. Santiago, E. Arbeille, and G. J. Bashaw, ”Robo2 acts in trans to inhibit Slit-Robo1 repulsion in pre-crossing commissural axons,” eLife, vol. 4, p. e08407, 2015/07/17 2015, doi: 10.7554/eLife.08407.

[14] A. Kerstetter-Fogle, H. Peggy, J. Barnholtz-Sloan, M. Couce, and A. Sloan, ”CSIG-38. ROBO2 SIGNALING IN INVASION OF GLIOBLASTOMA,” Neuro-Oncology, vol. 20, no. suppl_6, pp. vi51–vi51, 2018, doi: 10.1093/neuonc/noy148.204.

[15] R. K. Gara, S. Kumari, A. Ganju, M. M. Yallapu, M. Jaggi, and S. C. Chauhan, “Slit/Robo pathway: a promising therapeutic target for cancer,” (in eng), Drug Discov Today, vol. 20, no. 1, pp. 156–64, Jan 2015, doi: 10.1016/j.drudis.2014.09.008.

[16] L. H. Geraldo et al., “Monoclonal antibodies that block Roundabout 1 and 2 signaling target pathological ocular neovascularization through myeloid cells,” (in eng), Sci Transl Med, vol. 16, no. 774, p. eadn8388, Nov 20 2024, doi: 10.1126/scitranslmed.adn8388.

[17] S. A. Abdel-Rahman and M. T. Gabr, ”Optimization and development of a high-throughput TR-FRET screening assay for SLIT2/ROBO1 interaction,” SLAS Discovery, vol. 34, p. 100240, 2025/07/01/ 2025, doi: 10.1016/j.slasd.2025.100240.

[18] D. Wu, Q. Chen, X. Chen, F. Han, Z. Chen, and Y. Wang, ”The blood–brain barrier: Structure, regulation and drug delivery,” Signal Transduction and Targeted Therapy, vol. 8, no. 1, p. 217, 2023/05/25 2023, doi: 10.1038/s41392-023-01481-w.

[19] S. A. Abdel-Rahman and M. Gabr, “Small Molecule Immunomodulators as Next-Generation Therapeutics for Glioblastoma,” (in eng), Cancers (Basel), vol. 16, no. 2, Jan 19 2024, doi: 10.3390/cancers16020435.

[20] S. Upadhyay, B. Kaur, and M. T. Gabr, “CD28 and ICOS in immune regulation: Structural insights and therapeutic targeting,” (in eng), Bioorg Med Chem Lett, vol. 127, p. 130310, Nov 1 2025, doi: 10.1016/j.bmcl.2025.130310.

[21] G. Yom-Tov, R. Barak, O. Matalon, M. Barda-Saad, J. Guez-Haddad, and Y. Opatowsky, ”Robo Ig4 Is a Dimerization Domain,” Journal of Molecular Biology, vol. 429, no. 23, pp. 3606–3616, 2017/11/24/ 2017, doi: 10.1016/j.jmb.2017.10.002.

[22] L. Jendele, R. Krivak, P. Skoda, M. Novotny, and D. Hoksza, ”PrankWeb: a web server for ligand binding site prediction and visualization,” Nucleic Acids Research, vol. 47, no. W1, pp. W345–W349, 2019, doi: 10.1093/nar/gkz424.

[23] S. Upadhyay, V. Talagayev, S. Cho, G. Wolber, and M. Gabr, ”Structure-based virtual screening identifies potent CD28 inhibitors that suppress T cell co-stimulation in cellular and mucosal models,” European Journal of Medicinal Chemistry, vol. 300, p. 118194, 2025/12/15/ 2025, doi: 10.1016/j.ejmech.2025.118194.

[24] N. García-Vázquez and M. T. Gabr, ”Orthogonal temperature-related intensity change (TRIC) and TR-FRET as a high-throughput screening platform for the discovery of SLIT2 binders: A proof-of-concept approach,” SLAS Discovery, vol. 35, p. 100264, 2025/09/01/ 2025, doi: 10.1016/j.slasd.2025.100264.

[25] S. Upadhyay, S. Cho, H. Nada, and M. Gabr, “Discovery of CD28-Targeted Small Molecule Inhibitors of T Cell Co-stimulation Using Affinity Selection-Mass Spectrometry (AS-MS) and Ex Vivo Validation,” (in eng), bioRxiv, Aug 2 2025, doi: 10.1101/2025.07.31.667814.

[26] B. Kaur, H. Nada, L. Zhang, and M. T. Gabr, ”Lead optimization of a CHI3L1 inhibitor for Glioblastoma: Enhanced target engagement, pharmacokinetics, and efficacy in 3D spheroid models,” European Journal of Medicinal Chemistry, vol. 297, p. 117924, 2025/11/05/ 2025, doi: 10.1016/j.ejmech.2025.117924.

[27] J. Tang, N. Karbhari, and J. L. Campian, ”Therapeutic Targets in Glioblastoma: Molecular Pathways, Emerging Strategies, and Future Directions,” (in eng), Cells, vol. 14, no. 7, Mar 26 2025, doi: 10.3390/cells14070494.

[28] L. H. Geraldo et al., “SLIT2/ROBO signaling in tumor-associated microglia and macrophages drives glioblastoma immunosuppression and vascular dysmorphia,” (in eng), J Clin Invest, vol. 131, no. 16, Aug 16 2021, doi: 10.1172/jci141083.

[29] L. Calvo-Barreiro, S. Upadhyay, and M. T. Gabr, ”Temperature-related intensity change (TRIC)-based high-throughput screening enables the discovery of small molecule CD28 binders,” SLAS Discovery, vol. 35, p. 100256, 2025/09/01/ 2025, doi: 10.1016/j.slasd.2025.100256.

[30] S. Upadhyay, H. Nada, S. Cho, and M. T. Gabr, ”Overcoming the undruggable barrier: Structure-guided discovery of a potent small molecule CD28 antagonist with translational potential,” Biomedicine & Pharmacotherapy, vol. 194, p. 118937, 2026/01/01/ 2026, doi: 10.1016/j.biopha.2025.118937.

